# Contrasting contribution of embryo- and postnatally-derived brain-resident macrophages in sustaining sleep-wake circuitry

**DOI:** 10.1101/2024.01.22.576653

**Authors:** Ali Seifinejad, Mojtaba Bandarabadi, Meriem Haddar, Saskia Wundt, Mehdi Tafti, Anne Vassalli, Abbas Khani, Gianni Monaco

**Author notes:** Department of Biomedical Sciences, University of Lausanne, Lausanne, Switzerland (Ali Seifinejad). Correspondence to:* Dr. Ali Seifinejad or Dr. Gianni Monaco.

## Abstract

Sleep is a complex behavior regulated by various brain cell types. However, the roles of brain-resident macrophages, including microglia and CNS-associated macrophages (CAMs), particularly those derived postnatally, in sleep regulation remain poorly understood. Here, we investigated the effects of natural (embryo-derived) and repopulated (postnatally derived) brain-resident macrophages on the regulation of vigilance states. We found that depletion in embryonically-derived brain macrophages caused increased sleep in the active period, but reduced its quality, reflected in reduced power of brain sleep oscillations. This was observed both for the Non-REM and REM sleep stages. Subsequent repopulation by postnatal brain macrophages unexpectedly failed to reestablish normal sleep-wake patterns and additionally induced sleep fragmentation. Furthermore, brain macrophage depletion caused excitatory-inhibitory synaptic imbalance, which was resistant to repopulation, and led to increased inhibitory synapses. At the metabolite level, the distinct metabolite profile induced by brain macrophage depletion largely returned to normal after repopulation. Our findings suggest a so far largely unknown interaction between brain-resident macrophages and sleep and emphasizes striking functional differences between embryonic and postnatally-derived brain macrophages, paving the way to future exploration of the role of brain macrophages of different origin in sleep disorders and synaptic connectivity.

## Introduction

Sleep is a complex behavior essential for the normal functioning of animals, consistently linked to various diseases when compromised^1^. Sleep regulation involves circadian and homeostatic processes, determining respectively sleep timing and sleep need ^2^.

Sleep oscillatory patterns rely on the intricate connectivity of the central nervous system (CNS) at synaptic and neurotransmitter levels. Various cell types, primarily neuronal populations located in the hypothalamus, brainstem (BS), and basal forebrain (BF), interact to finely regulate sleep. The expression of synaptic proteins increases during wakefulness and sleep loss, and decreases during sleep^3^, and the postsynaptic protein HOMER1A is a well-established core brain molecular correlate of sleep loss ^4^. Additionally, the release of key neuropeptides and neurotransmitters such as HCRT (also called Orexin), serotonin (5HT), noradrenaline (NA), and acetylcholine across different brain regions actively modulates sleep-wake cycles^5^. This intricate interplay manifests in electroencephalographically (EEG) measurable readouts distinguishing wakefulness, non-rapid eye movement sleep (NREMS), and rapid eye movement sleep (REMS), characterized by distinct brain oscillatory activities such as delta waves (0.5-4 Hz) for NREMS and theta waves (6-9 Hz) for REMS in rodents. However, the contribution of non-neuronal cell types, such as brain-resident macrophages, to brain oscillations and sleep architecture remains largely unexplored.

Brain-resident macrophages comprising parenchymal microglia and CAMs ^6–8^, play significant roles in pathology and protecting the CNS from damage ^9^. They participate in a range of physiological functions including synaptic pruning, neurogenesis, immune surveillance ^10–12^, and are derived from prenatal sources known as erythromyeloid precursors that engraft into the developing CNS ^13,14^, facilitating the establishment of neuronal networks especially in the first postnatal weeks ^10^. If embryo-derived brain-resident macrophage cells are depleted in adulthood, they repopulate from surviving endogenous cells ^15^. D*e novo* establishment of micoglial networks occurs by surviving cells through clonal expansion ^16^ and these postnatal-derived microglial cells exhibit a different transcriptional signature ^17^. However, whether adulthood-derived microglia exhibit functional differences relative to orginal embryo-derived populations is hotly disputed.

There is evidence that immune system activation, such as during illness, results in increased sleepiness, suggesting an interaction between the immune system and the sleep-wake cycle ^18,19^. Brain-resident macrophages, being the closest immune components to sleep-regulating centers in the brain, might play a crucial role in this interaction. While some studies have demonstrated a clear influence of sleep loss on embryo-derived microglia ^20–22^, the opposite relationship and how the brain-resident macrophages modulate sleep/wake cycle and vigilance states remain largely unexplored. Therefore, it is essential to thoroughly investigate the extent to which brain-resident macrophages influence sleep states and brain activity during these states. EEG-recorded brain activity will therefore be an important indicator of the effects of the immune system on brain oscillations. Additionally, the influence of brain macrophages on sleep-regulating neurotransmitters, and synaptic connectivity warrants further exploration. Finally, it remains unclear whether adulthood-derived brain-resident macrophages are capable of re-establishing neural circuitries in the same manner as prenatal cells, including synaptic plasticity and neurochemical balance, which are crucial for normal sleep regulation. Here, we undertook to address these questions through a sound experimental design and in-depth analyses. Using a comprehensive characterization of EEG signals recorded from the mice brain, we first explored whether the absence of embryo-derived brain-resident macrophages influences sleep-related brain oscillatory patterns. Next, we studied whether repopulated postnatal macrophages can normalize these patterns. Finally, we investigated how the absence and re-presence of microglia and CAMs affects brain network connectivity and neurochemistry. Our data show functional differences between embryo-derived and postnatally adulthood-derived brain-resident macrophages.

## Results

### Repopulated postnatally-derived brain-resident macrophages are deficient in rebuilding normal NREMS circuitry

We examined to what extent embryo-derived brain-resident macrophages contribute to sleep regulation and whether postnatal-derived macrophages mirror the same functionality. To address this, we depleted microglia from the brains of mice for two weeks using colony-stimulating factor 1 receptor (CSF1R) inhibition via PLX5622 ^23^, and then allowed them to repopulate for four weeks (Fig. 1A). CSF1 receptor is vital for microglia survival. Antagonizing it severely depletes microglia within three days, and upon the termination of treatment, repopulation can be readily observed within a few days^17^. We observed substantial microglia depletion (>86%) after two weeks of PLX5622 treatment, and complete repopulation four weeks after withdrawal (Fig. 1B, C).

**Figure 1.**
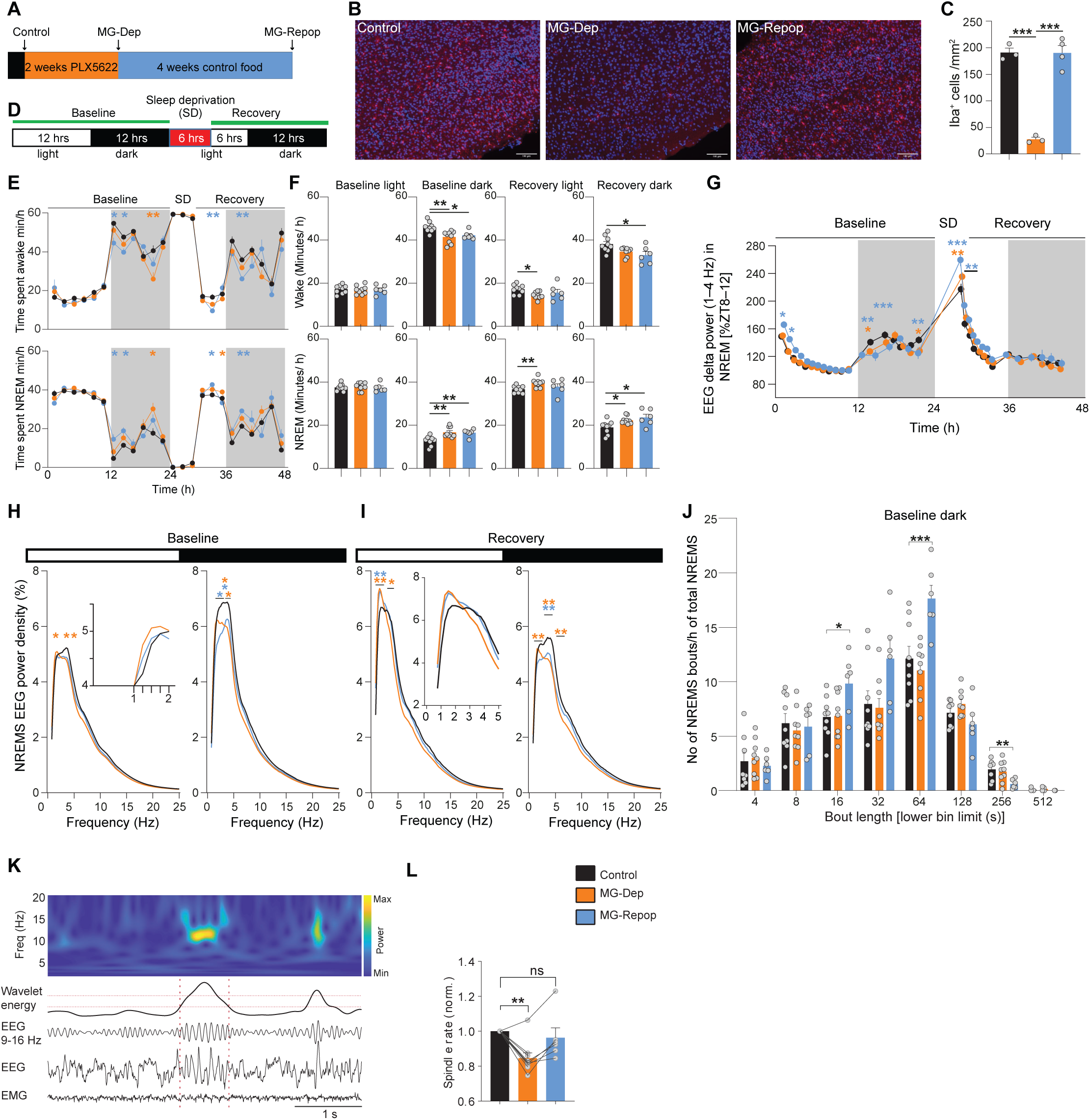
Essential role of MG in regulation of NREMS and associated EEG oscillations. **(A)** Schematic illustration of the experimental procedure conducted on adult mice to assess the brain’s electrical activity using EEG. Recordings prior to PLX5622 drug administration served as controls. Mice were then fed PLX5622-containing food during 2 weeks. During the final 3 days of this 2-week period, a 2^nd^ series of recordings was conducted. Following PLX5622 treatment, mice were provided control food for 4 weeks. After 4 weeks under normal food, a 3^rd^ recording was performed. All 3 recordings were performed on the same mice. (**B, C**) PLX5622 substantially depleted the microglia from the brain within 2 weeks (compare control and MG-Dep). Return to normal food repopulated microglia into the brain after 4 weeks (compare MG-Dep and MG-Repop). (**D**) Recording paradigm. The first 2 days of recordings served as baseline, averaged as one day. The 2^nd^ day consisted of 6 h SD followed by 18 h of recovery. (**E**) Time-course of vigilance states; Wakefulness (top) and NREMS (bottom). Compared to controls (black datapoints), MG-Dep (orange datapoints) and –Repop (blue datapoints) mice show a large increase in NREMS duration during the dark period both at baseline and during recovery (n = 9 for control and MG-Depleted conditions and n = 6 for MG-Repopulated condition, 2-way ANOVA, followed by Dunnett’s test, *P < 0.05; **P < 0.01; ***P < 0.001, orange and blue stars, significant differences between control and MG-Dep and MG-Repop mice, respectively). Data points are shown in minutes per hour (min/h) and represent the average of 2 h. (**F**) From left to right: Time in wakefulness (top) and NREMS (bottom) during baseline light, dark and recovery light and dark periods (Mixed-effects analysis, followed by Dunnett’s test, *P < 0.05; **P < 0.01; ***P < 0.001). (**G**) Time-course of EEG delta power (1–4 Hz) during the 3-day recording. EEG delta power is a proxy of sleep need. Both MG-Dep and MG-Repop mice show deficiency in building up sleep need during the dark period (2-way ANOVA, followed by Dunnett’s test, *P < 0.05; **P < 0.01; ***P < 0.001, orange and blue stars, significant differences between control and MG-Dep and MG-Repop mice, respectively). (**H**) EEG power spectra of experimental mice in NREMS during baseline light (left) and dark (right) periods. EEG power is expressed as % of baseline power of control condition. Insets show amplification of specific EEG frequencies (2-way ANOVA, followed by Dunnett’s test, *P < 0.05; **P < 0.01; ***P < 0.001, orange and blue stars indicate significant differences between control and MG-Depleted and MG-Repopulated mice, respectively). (**I**) EEG power spectra of experimental mice in NREMS during recovery light (left) and dark (right) periods. (**J**) Distribution of NREMS bout durations during baseline dark period across the experimental mice. Sleep bout duration is represented on the x-axis (1-way ANOVA, followed by Tukey test). (**K**) (Top) Representative time-frequency heatmap of spindles. (Bottom) Representative EEG/EMG signals of a detected spindle. Dashed horizontal lines indicate upper and lower thresholds used to detect spindles with wavelet energy within 9-16 Hz, and dashed vertical lines indicate the start and end of the detected spindle. (**L**) Spindle rate during NREMS episodes in different conditions. Spindle rate significantly decreased in MG-depleted mice (mixed-effects analysis, followed by Dunnett’s test, *P < 0.05; **P < 0.01; ***P < 0.001)).

We recorded EEG signals before depletion (control), after depletion and after repopulation (Fig. 1A). Since CSF1 inhibition also depletes CAMs ^24^, the effects we report here are due to the depletion of both cell types, which we refer to collectively as MG. Our experimental conditions therefore included control, MG-Dep (microglia/CAM depleted), and MG-Repop (microglia/CAM repopulation). To evaluate the distribution of vigilance states, we employed our standard sleep/wake phenotyping protocol ^25^, with two days of baseline (which were averaged and presented as one day in the figures) and in the third day mice underwent 6 h of SD (ZT0-ZT6) and 18 h of recovery (Fig. 1D). In baseline conditions, all experimental mice demonstrated the characteristic cycling behavior, with higher levels of wakefulness during the dark period and higher levels of NREMS during the light period and exhibited the increase in NREMS amounts in response to SD (Fig. 1E, F). However, a significant NREMS increase was observed in the baseline dark period when comparing the MG-Dep to the control condition. Interestingly, the repopulation of MG did not normalize this effect (Control: 12.75±0.72 min, MG-Dep: 16.78±0.77 min, MG-Repop: 16.54±0.70 min, mean ± SEM) (Fig. 1E, F, bottom panels). Reciprocally, there was a notable reduction in wakefulness during the dark period (Fig. 1E, F, top panels).

To gain insight into sleep homeostatic regulation in different mouse groups and determine whether increased sleep is due to increased need for sleep, we analyzed the time course of NREMS EEG delta power (1-4 Hz, a recognized quantitative readout of sleep need) across our 3-day recordings. All mice groups showed a typical decline in delta power during the light period and a strong rebound after SD, indicating homeostatic regulation of sleep need in all groups (Fig. 1G). However, the buildup in delta power during the baseline dark period was significantly lower in MG-Dep and MG-Repop mice compared to WT mice (Fig. 1G). This attenuation was more pronounced in MG-Repop mice and may be partly related to the increased amount of sleep during the baseline dark period (Fig. 1E, lower panel). In contrast, when mice were exposed to enforced wakefulness (SD), the EEG delta power increase was significantly more pronounced in MG-Dep and MG-Repop than in control conditions, and again this effect was more extensive in MG-Repop than in MG-Dep (Fig. 1G). Altogether, these data indicate impaired buildup of homeostatic sleep need during the waking period.

Spectral analysis of NREMS revealed that MG-Dep mice exhibited higher slow-delta (1-2 Hz) activity compared to control mice in baseline light period (Fig. 1H, left). This frequency band appears particularly affected also in MG-Repop mice, as it was significantly increased in the recovery light period following SD (Fig. 1I, left). The fast-delta (3-4 Hz) frequency range was in contrast blunted during both baseline and recovery light periods only in the MG-Dep condition (Fig. 1H, I, left). During the baseline dark period, large delta power blunting was observed in both MG-Repop (1.5-3.5 Hz) and MG-Dep (3-5.5 Hz) conditions. These deficits remained nearly the same during the recovery dark period, indicating that NREMS of both experimental mice appears to be shallower (Fig. 1H, I, right). Collectively, it can be concluded that the presence of MG is necessary for the precise regulation of fast and slow-delta power as a proxy of the sleep homeostatic process. MG repopulation not only fails to restore NREMS regulation but worsens it.

To determine whether increased NREMS duration is due to enhanced state initiation (increased number of NREMS episodes) and/or to enhanced state maintenance (increased NREMS episode duration), we conducted a sleep fragmentation analysis. We found that while MG-Dep mice have similar sleep bout duration patterns as the control condition, MG-Repop mice showed a larger number of shorter NREMS bouts (16 seconds [Control: 6.7±0.63, MG-Dep: 6.8±0.7, MG-Repop: 9.84±1.12]) and 64 seconds [Control: 12.12±1.15, MG-Dep: 11.02±0.81, MG-Repop: 17.63±1.2]) and fewer longer NREMS bouts (>256 seconds [Control: 1.96±0.28, MG-Dep: 1.74±0.33, MG-Repop: 0.62±0.17]), suggesting a strong destabilization of NREMS in the MG-Repop brain (Fig. 1J).

During NREMS, brief bursts of brain activity in the 9-16 Hz frequency range, called spindles, are generated by thalamo-cortical interactions and play roles in processes such as learning, memory, and cognition ^26^. To determine if MG depletion and repopulation influence spindle generation, we quantified spindle rate during NREMS episodes and found a significantly reduced spindle rate in MG-Dep mice compared to the control condition (Fig. 1K, L; see “Methods”). In contrast to other features of NREMS, MG repopulation was found to normalize the occurrence rate of spindle events, indicating that MG repopulation partially rescues the circuit deficits seen after MG depletion.

Altogether, these data suggest that embryo-derived MG are an integral part of NREMS establishment and maintenance, and repopulated postnatal MG largely fail to fulfill the normal NREMS regulatory functions of embryo-derived MG.

### Depletion of brain-resident macrophages prolongs REMS and degrades its quality

Different neural pathways and neurotransmitters regulate REMS compared to NREMS. However, the role of MG in the regulation of REMS is not well understood. We quantified REMS amount across the dark/light periods and found that MG-Dep mice display more REMS compared to the control and MG-Repop conditions during the baseline dark period (Control: 0.67±0.1, MG-Dep: 1.17±0.21 min/h, mean ± SEM). However, in contrast to NREMS, MG repopulation corrected the abnormal expression of REMS (MG-Repop: 0.61±0.13 min/h) (Fig. 2A, B). To investigate if the increase in REMS is due to the consolidation of this state, we performed fragmentation analysis and found that the number of shorter REMS bouts (8 s) is largely decreased (Control: 8.7±2.05, MG-Dep: 2.8±1.2, MG-Repop: 8.8±3.3) and longer bouts (>2 min) are increased (Control: 1.5±0.8, MG-Dep: 6.1±1.1, MG-Repop: 4.71±1.4, mean ± SEM) (Fig. 2C) in MG-Dep mice, suggesting stabilization of REMS in the absence of MG. Theta oscillations dominate the EEG signal during REMS, where theta phase dynamically modulates gamma amplitude in hippocampal and cortical networks ^27^. We next investigated the contribution of MG in theta oscillations and theta-gamma interactions. Spectral analysis of REMS revealed a significant dampening of theta (6-8 Hz) power in MG-Dep mice during the baseline dark period (Fig. 2D), which was restored after MG repopulation. We also found that the theta-gamma coupling is significantly reduced in the MG-Dep condition compared to the control. Although MG repopulation attempts to bring it back to the normal level, it fails significantly (Fig. 2E, F).

**Figure 2.**
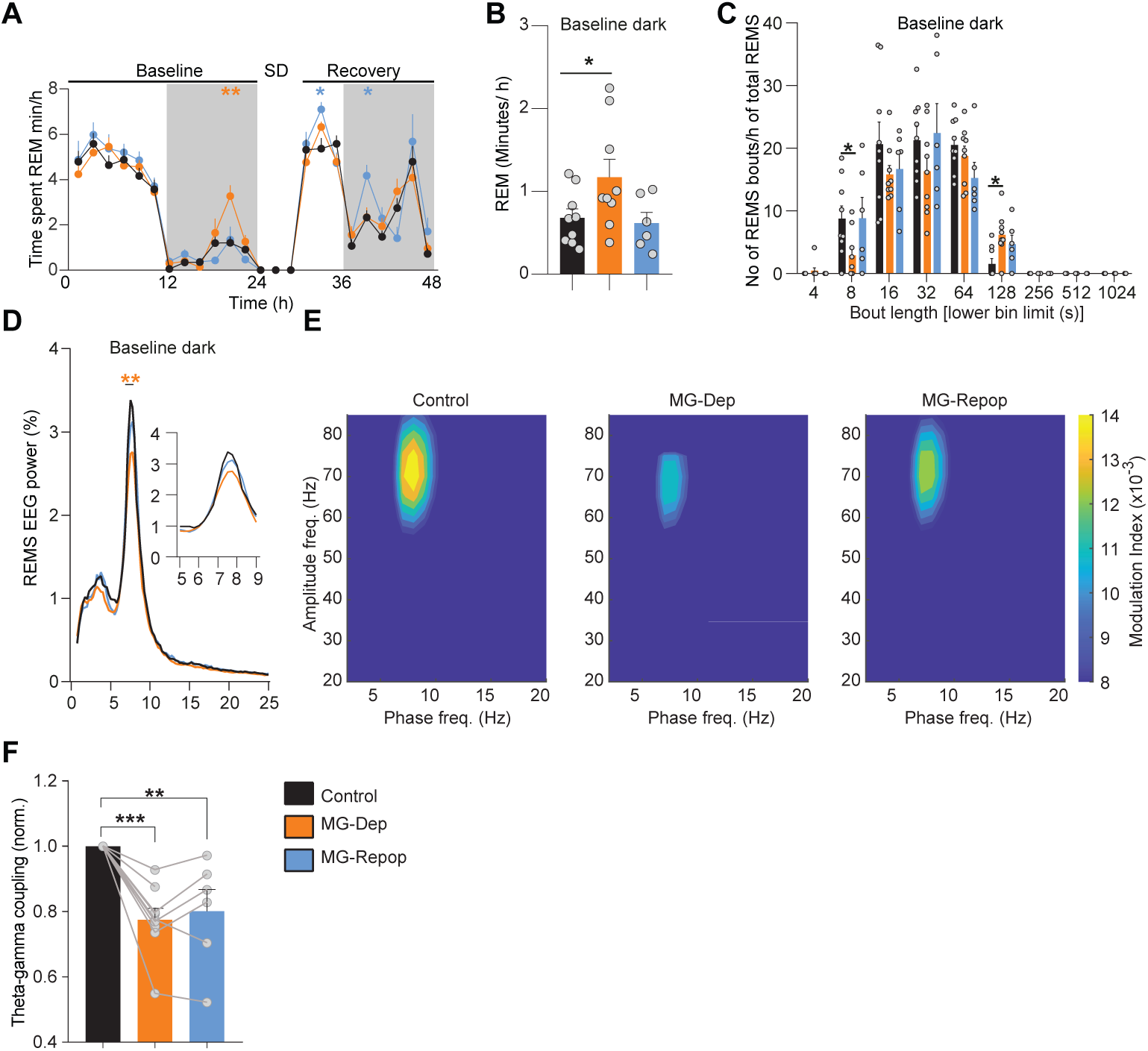
REMS regulation largely depends on the presence of the MG. (**A**) Time-course of REMS duration in control (black), MG-Dep (orange) and MG-Repop (blue) mouse groups. MG-Dep mice exhibit a large increase in REMS time during the baseline dark period (n = 9 for control and MG-Dep conditions and n = 6 for MG-Repop condition, 2-way ANOVA, followed by Dunnett’s test, *P < 0.05; **P < 0.01; ***P < 0.001, orange and blue stars represent significant differences between control and MG-Dep and MG-Repop mice, respectively). Data points are shown in minutes per hour (min/h) and represent the average of 2 h. (**B**) Time in REMS during baseline dark period (mixed-effects analysis, followed by Dunnett’s test, *P < 0.05; **P < 0.01; ***P < 0.001). (**C**) Distribution of REMs bout durations during baseline dark period across 3 mouse groups. Sleep bout duration is presented as x-axis (1-way ANOVA, followed by Tukey test). (**D**) EEG power spectra of all experimental mice in REMS during baseline dark period. Insets show amplification for specific EEG frequencies (2-way ANOVA, followed by Dunnett’s test, *P < 0.05; **P < 0.01; ***P < 0.001, orange and blue stars are significant differences between control mice and MG-Dep and MG-Repop mice respectively). (**E**) Comodulogram graphs show the modulation index for a wide range of frequency pairs, obtained from 12-h recordings of one animal in different conditions. (**F**) Normalized theta-gamma coupling across 3 experimental conditions. Theta-gamma coupling is significantly decreased in MG-depleted and -repopulated conditions (mixed-effects analysis, followed by Dunnett’s test, *P < 0.05; **P < 0.01; ***P < 0.001)).

Altogether, these findings suggest that the presence of MG is necessary for the normal expression of REMS.

### Brain-resident macrophages contribute to the maintenance of active wakefulness

The quality of prior wakefulness contributes to the expression of both NREMS and REMS. Along these lines, a specific sub-state of prior wakefulness was recently proposed to drive NREMS need ^28^, and some aspects of wakefulness may functionally substitute for REMS ^29^. To investigate how wakefulness is expressed in our experimental conditions, we performed a dynamic analysis of the full waking EEG spectrum across three recording days. This global analysis revealed differential expression of the higher waking frequency bands during light-dark cycles across three days, and most notably during enforced wakefulness (SD). As suggested in Figure 3 (Fig. 3A), both MG-Dep and MG-Repop mice exhibit a large dampening of the lower gamma (≈45-55 Hz) band during the baseline dark period, which is exacerbated during the recovery dark period. Additionally, the MG-Dep mice exhibit a large blunting of the higher gamma band (≈70-90 Hz) during SD, which partially normalizes after MG repopulation. To better quantify the lower frequency bands, we performed PSD analysis and found that MG-Dep mice exhibit a large increase in slow delta (1.35-1.75 Hz) activity compared to the control mice (Fig. 3B).

**Figure 3.**
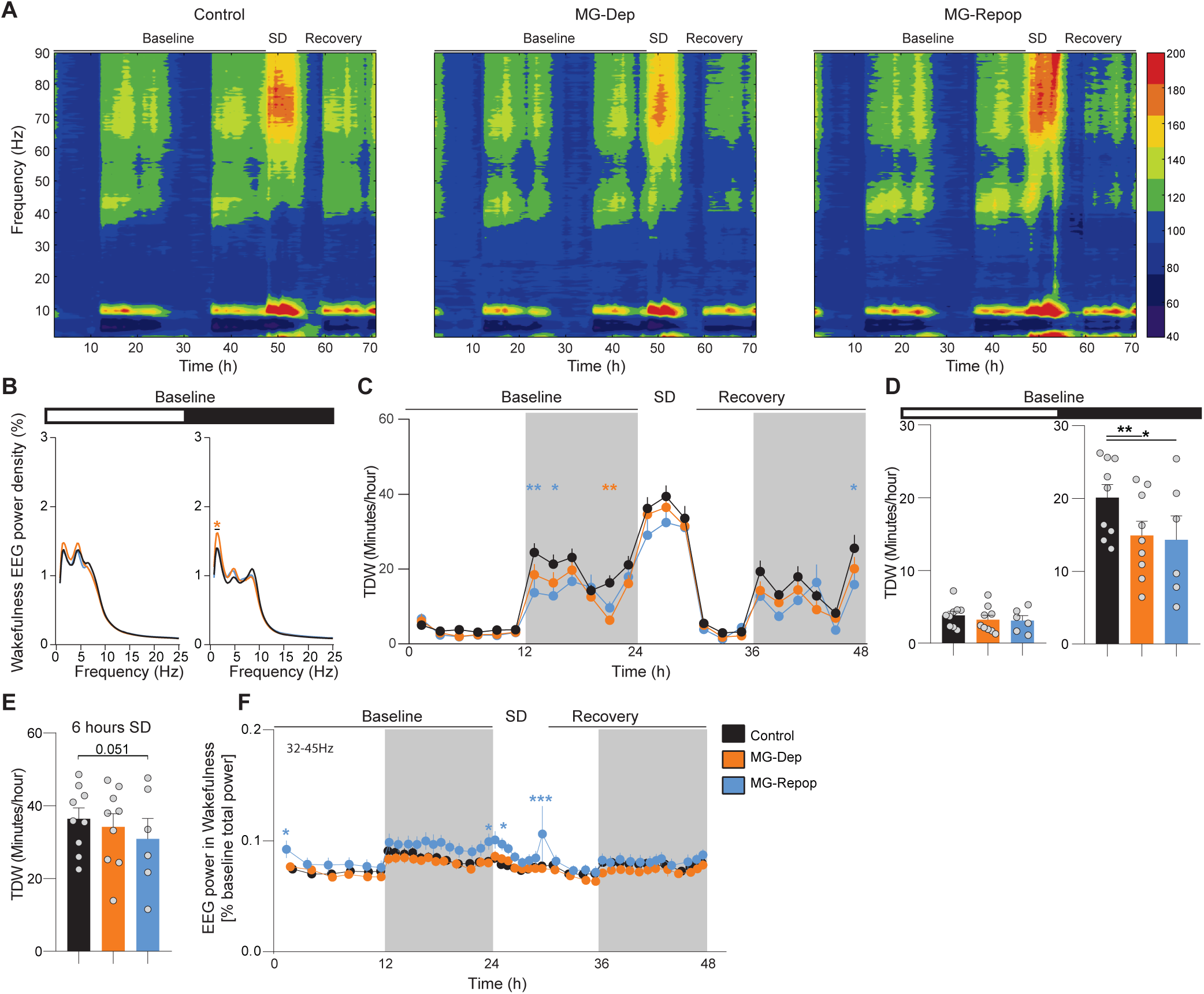
MG Repopulation fails to normalize the waking state deficits induced by MG depletion. (**A**) Time x frequency x power heatmap representations of the waking EEG of control, MG-Dep and MG-Repop mice across the 3 recorded days (48 h baseline, 6 h sleep deprivation and 18 h recovery) show large changes in higher frequency bands. (**B**) EEG power spectra of experimental mice in wakefulness during baseline light and dark periods (2-way ANOVA, followed by Dunnett’s test, *P < 0.05; **P < 0.01; ***P < 0.001, orange star are significant differences between control mice and MG-Dep mice). (**C**) Time-course of TDW amounts indicates decreased TDW time during the dark period in MG-Dep and MG-Rep. Datapoints are shown in min/h and represent the average of 2 h (baseline days 1 and 2 are averaged), (2-way ANOVA followed by Dunnett’s test, orange and blue stars are significant differences between control and MG-Dep and MG-Repop mice, respectively). (**D**) Quantification of time in TDW (min/h) during baseline light and dark period (mixed-effects analysis, followed by Dunnett’s test, *P < 0.05; **P < 0.01; ***P < 0.001). (**E**) Time in TDW during SD. (**F**) Waking EEG slow-gamma (32-45 Hz) power dynamics during the 3-day recording (baseline days 1 and 2 are averaged) (2-way ANOVA, followed by Dunnett’s test).

Mice spend a portion of their waking time engaged in exploratory and motivated behaviors, such as running, nest building, or drinking, often referred to in rodents as “active wakefulness”. These behaviors are associated with the EEG signature called “theta-dominated wakefulness” (TDW 6.0-9.5 Hz), which has been suggested to be the principal driver of the sleep homeostat^28^. We quantified TDW in our mice and found a large decrease in its amount in both MG-Dep and MG-Repop mice during the baseline dark period (Fig. 3C, D). This reduction was more pronounced in MG-Repop mice, which was also seen during SD (Fig. 3C, E). TDW is associated with heightened gamma activity. We analyzed the time course of gamma activity during wakefulness and found an increase in slow (32-45Hz) frequency bands in the MG-Repop condition compared to the controls (Fig. 3F). The increase in these frequency bands in MG-Repop mice is also apparent on time-frequency heat maps in Figure 3A.

Overall, these data suggest a significant imbalance in the theta/gamma-rich waking state in both the MG-Dep and MG-Repop conditions, and appearance of heightened gamma activity after MG repopulation.

### Disrupted balance of synaptic signaling upon MG depletion and repopulation

Sleep is associated with changes in synaptic plasticity ^30^ and microglia play a substantial role in synaptic functions, which are dependent on neuronal activity and vigilance states (VS)^31^. To investigate whether MG depletion and repopulation induce changes in brain network connectivity at the synaptic level, we analyzed a subset of excitatory and inhibitory synaptic density proteins that are involved in sleep regulation, (e.g., the core brain molecular correlate of sleep loss, HOMER1) in the brains of our experimental mice.

Using an ImageJ macro, we counted the number of synapses on cortical and hypothalamic neurons (Fig. 4A, B) and found that the number of excitatory VGLUT1-HOMER1 synapses on NEUN^+^ cortical neurons was significantly reduced in both MG-Dep and MG-Repop conditions. This reduction was more noticeable in MG-Repop mice (Fig. 4C). Interestingly, this effect was regulated at the postsynaptic level (HOMER1-associated, Fig. 4D) rather than the presynaptic level (VGLUT1-associated, Fig. 4E). Recently, cortical PV neurons have been proposed to be responsible for homeostatic sleep regulation^32^. We found that the number of parvalbumin (PV)-originated^33^ Synaptotagmin-2-Gephyrin (Syt2-Geph) inhibitory synapses on the cortical NEUN^+^ neurons was significantly increased under the MG-Repop condition (Fig. 4F, G). This increase could possibly be due to an increase in the activity of PV neurons, as we found that the number of VGLUT1-HOMER1 excitatory synapses on PV neurons was significantly increased under both MG-Dep and MG-Repop conditions (Fig. 4H, I). We additionally investigated the changes in the count of VGLUT1-HOMER1 synapses on representative hypothalamic neurons (HCRT) and found a significant decrease in the number of VGLUT1-HOMER1 synapses on HCRT neurons in both MG-Dep and MG-Repop conditions (Fig. 4J, K).

**Figure 4.**
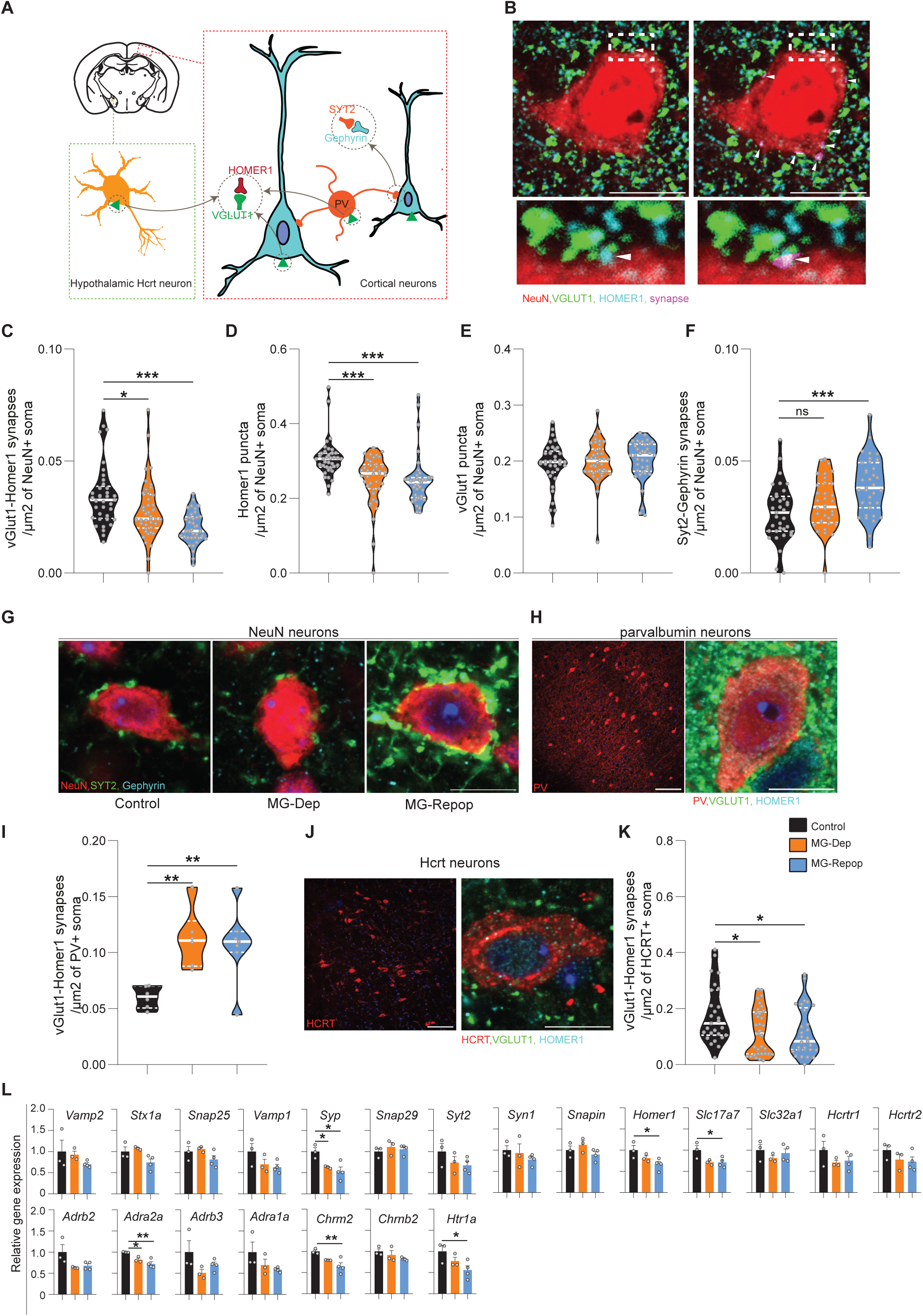
Large synaptic imbalance after MG depletion and repopulation. (**A**) Schematic illustration of the cell types for which the number of synapses was counted. NEUN (blue) and PV (orange) positive neurons in the cortex and HCRT neurons in the hypothalamus. The number of co-localized excitatory VGLUT1-HOMER1 synaptic proteins on NEUN, PV and HCRT positive neurons were counted. Additionally, the number of co-localized inhibitory SYT2-Gephyrin synaptic proteins on NEUN neurons were counted. (**B**) Representative image depicting pre, postsynaptic proteins and their colocalization in a NEUN positive neuron. Synapses (arrowheads) are marked with pink color using ImageJ macro (right). Scale bar: 10µm. (**C**) Number of VGLUT1-HOMER1 synapses/μm^2^ of NEUN^+^ soma. N=342 cells from 12 mice for control, 324 cells from 11 mice for MG-Dep, and 357 cells from 12 mice for MG-Repop conditions. The data show a reduced number of these synapses in both MG-Dep and MG-Repop conditions (**D**) Number of HOMER1 puncta/μm^2^ of NEUN^+^ soma. (**E**) Number of VGLUT1 puncta/μm^2^ of NEUN+ soma. (**F**) Number of SYT2-Gephyrin synapses/μm^2^ of NEUN^+^ soma. N=299 cells from 12 mice for control, 270 cells from 11 mice for MG-Dep, and 306 cells from 12 mice for MG-Repop conditions. (**G**) Representative images depicting SYT2-Gephyrin synapses in NEUN^+^ cortical neuron. Large increase in these inhibitory synapses is observed in the MG-Repop condition. Scale bar: 10 µm. (**H**) Distribution of PV^+^ neurons (red) in the cortex. Scale bar: 100 µm (left), 10 µm (right). (**I**) Number of vGLUT1-HOMER1 synapses/μm^2^ of PV^+^ soma. N=29 cells from 5 mice for control, 29 cells from 5 mice for MG-Dep, and 23 cells from 4 mice for MG-Repop conditions. (**J**) Left, distribution of HCRT^+^ neurons (red) in the lateral hypothalamus. Right, a HCRT^+^ neuron stained with VGLUT1 (green) and HOMER1 (Cyan). Scale bar: 100 and 10 µm. (**K**) Number of VGLUT1-HOMER1 synapses/μm^2^ of HCRT+ soma. N=70 cells from 3 mice for control, 73 cells from 3 mice for MG-Dep, and 77 cells from 4 mice for MG-Repop conditions. (**L**) Gene expression of synaptic proteins and neurotransmitter receptors at RNA level following MG depletion and repopulation. N=3 for control and MG-Dep and n=4 for MG-Repop conditions. 1-way ANOVA, followed by Tukey test, *P < 0.05; **P < 0.01; ***P < 0.001.

To further investigate changes in network connectivity in the brains of MG-Dep and MG-Repop mice, we analyzed additional synaptic markers and sleep-specific neurotransmitter receptors (e.g., receptors of monoaminergic systems and neuropeptides) on RNA extracted from cortical samples of these mice. Our bulk analysis revealed that some general synaptic markers, such as *Synaptophysin I* (*Syp*), are largely downregulated in both experimental mice, while *Slc17a7* (encoding Vesicular glutamate transporter 1) and *Homer1* genes are significantly downregulated in the cortex of the MG-Repop condition. Among the specific neurotransmitter markers analyzed, the adrenergic receptor family (e.g. *Adrb2*, *Adra2a*, *Adrb3*, and *Adra1a*) genes showed a tendency to be less expressed in both MG-Dep and MG-Repop conditions, with a marked decrease in the expression of the *Adra2a* gene. Additionally, we observed a reduction in expression of *Cholinergic receptor muscarinic 2* (*Chrm2*) and the serotonergic receptor *5-Hydroxytryptamine receptor 1A* (*Htr1a*) genes, which mainly occur in the brain of MG-Repop mice (Fig. 4L).

Taken together, these data suggest that brain connectivity at the synaptic level is largely impaired following the depletion of brain-resident macrophages. Repopulating the brain with these macrophages not only failed to normalize connectivity but, in many cases analyzed, worsened it.

### Neurotransmitter release is largely restored after the repopulation of resident macrophages

Synapses, in addition to providing structural connections in the brain, play a crucial role in transmitting neurotransmitters essential for the function of complex neural systems. We therefore questioned whether the levels of essential neurotransmitters involved in sleep regulation are affected by the depletion and repopulation of brain-resident macrophages. We conducted targeted metabolomics to analyze the neurotransmitter content and metabolite reservoir of the cortex, target of most neurotransmitters and region where EEG signals are acquired. Assessment of neurotransmitter content was performed using chromatography with tandem mass spectrometry (LC-MS/MS) in the brain of control, as well as MG-Dep and MG-Repop mice. We found that pharmacological depletion of MG led to a distinct clustering of metabolites, as indicated by principal component analysis (PCA) (Fig. 5A). This suggests that MG plays an important role in maintaining the overall metabolic composition of the cortex. MG repopulation however restored the cortical submetabolome to a pattern similar to the one observed under control experimental conditions (Fig. 5A). Univariate analysis revealed a reduction in norepinephrine upon MG depletion, which was restored after repopulation (Fig. 5B). The concentrations of acetylcholine, choline, dopamine, and serotonin remained unchanged between the experimental conditions, with a slight tendency to decrease in MG-Dep condition.

**Figure 5.**
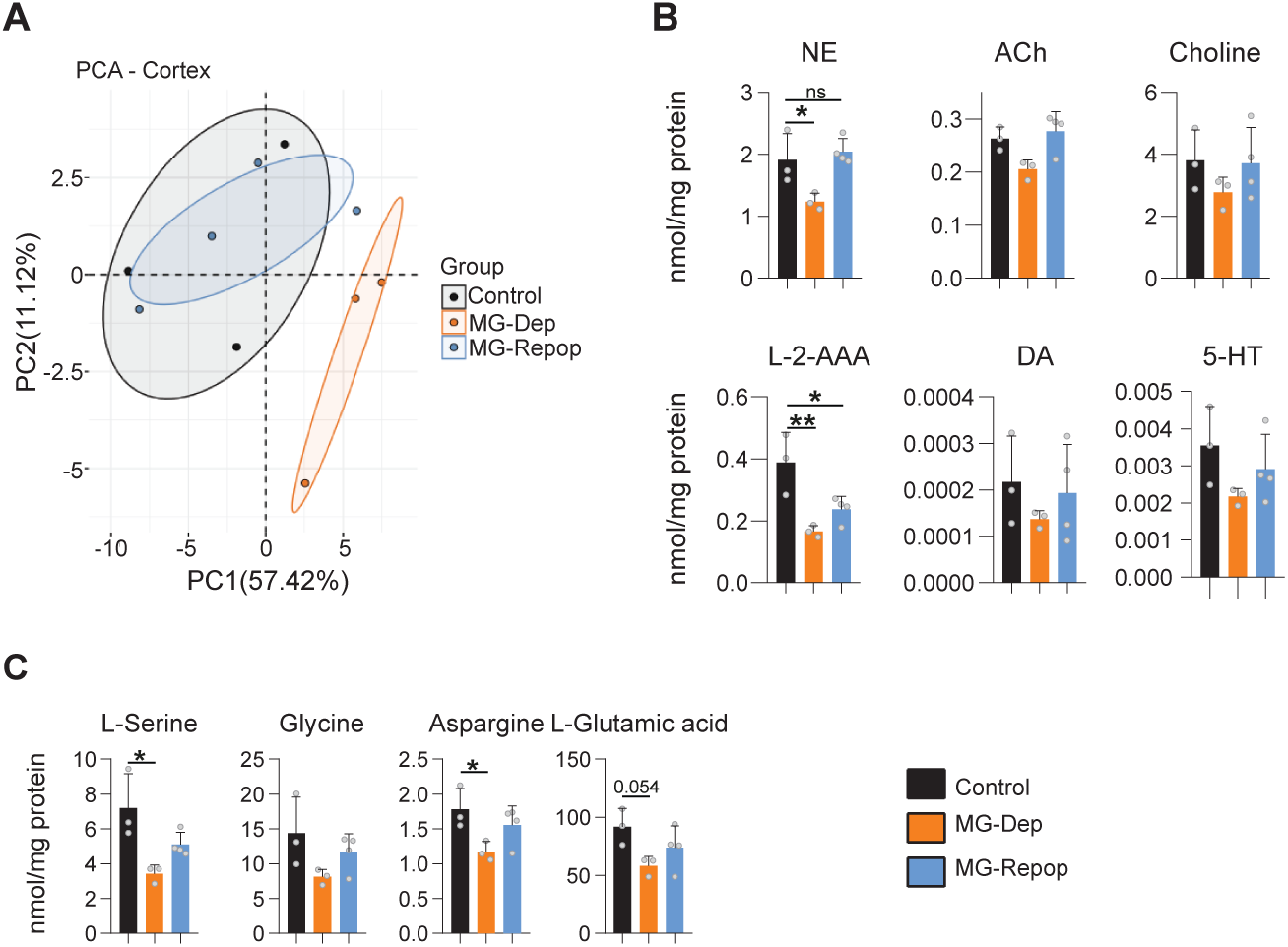
MG repopulation largely restores the metabolite changes induced by microglial depletion. (**A**) PCA of targeted metabolites in cortex isolated from control (n=3), MG depleted (n=3) and repopulated (n=4) mice. (**B**) LC-MS/MS assessment of the neurotransmitter content in the brain of control, MG-Dep and MG-Repop mice. (**C**) Downregulation of amino acids in the brain of experimental mice after MG depletion. 1-way ANOVA, followed by Tukey test, *P < 0.05; **P < 0.01; ***P < 0.001.

Furthermore, we observed that MG depletion led to a marked reduction of L-2-amino adipic acid (L-2-AAA), which was only partially recovered after repopulation (Fig. 5B). We also analyzed the precursors of neurotransmitters and found a significant decrease in the amino acids L-serine and asparagine under depleted condition, which could be restored upon MG repopulation (Fig. 5C). Similarly, there was a trend towards decreased glycine and L-glutamic acid content (Fig. 5C).

Taken together, our data suggest that while network connectivity is significantly impaired in MG-Dep condition and resistant to MG repopulation, the underlying substrates (neurotransmitters and metabolites) are largely restored after MG repopulation.

## Discussion

In this study, we investigated the extent to which embryo-derived brain-resident macrophages contribute to sleep regulation and whether postnatally derived ones serve a similar role. We demonstrate that MG depletion has a significant impact on the homeostatic regulation of sleep. The brain circuits most vulnerable to microglia depletion and repopulation, causing the changes in sleep EEG that we document, are still unknown. However, changes in slow-delta activity in NREM sleep, state stabilization and reduced theta power in REMS, as well as changes in waking slow-delta activity, were reported in mice with defects in hypothalamic HCRT and brainstem serotonergic and noradrenergic systems^25,28,34,35^. This is suggestive of functionally important local interactions between microglial populations and sleep regulating circuits, as mentioned above.

We showed that the changes in sleep patterns after MG depletion are not fully reversed after MG repopulation. Previous studies have shown that the brain global transcriptomes are little influenced by repopulated microglia ^17^. Furthermore, it has been suggested that after MG repopulation, neuroprotection is achieved through IL-6 induction by neurons^36^. Other studies reported full functionality of retinal microglia after repopulation, including continuous retinal surveillance, maintenance of synaptic structure, and normal behavioral and physiological responses to retinal injury ^37^. However, our data reveal that MG repopulation not only fails to completely restore various aspects of sleep regulation, but in some cases exacerbates altered parameters, such as fragmentation of NREMS and dysregulation of sleep need. This demonstrates striking funtional differences between microgial cells derived from pre-or postantal sources. We speculate that not all microglia-neuronal interactions are properly reestablished by adulthood-derived microglia after repopulation. Whether a longer repopulation period might correct the abnormalities observed in our analysis warrants further investigation. We observed a significant imbalance between excitatory and inhibitory synapses after depletion of embryo-derived microglia and after repopulation with adult microglia, with a notable increase in inhibitory synapses following repopulation. Studies in juvenile mice prevously reported that microglia depletion in the developing brain significantly increases the number of both excitatory (VGLUT1-HOMER1) and inhibitory (Syt2-Gephyrin) synapses on cortical neurons, an effect that persisted after repopulation until P30. However, in the adult (P60) brain, synapse density returned to normal control levels, suggesting that depleting microglia during development causes long-lasting, albeit not permanent, effects^38^. Our data, however, show that depleting microglia in adulthood, and then repopulating them, induces different results, with a reduction in excitatory synapses in both MG-Dep and MG-Repop conditions, and an increase in inhibitory synapses in MG-Repop conditions. In the developing brain, it has been suggested that MG-Dep impairs synaptic pruning processes and alters related gene expression^10,38^. Whether the same process or any specific pathways are induced by MG depletion and repopulation in the adult brain, such as aberrant reactivation of developmental programs^39^, remains to be found.

Furthermore, other studies have suggested that microglia play a role in restraining excessive neuronal activation that cannot be sufficiently suppressed by inhibitory neurons alone^40^. Our results suggest that embryo-derived microglia also control excessive activity of inhibitory PV neurons, but the reasons why this is not corrected with adulthood-derived microglia needs further investigation.

Deficient synaptic neurotransmission was proposed as a potential cause for abnormal sleep regulation^41^. We found here an imbalanced neurotransmitter and metabolite content of the brain in mice devoid of prenatal microglia. It appears that norepinephrine neurotransmission is particularly sensitive to MG deficiency, as both the levels of norepinephrine metabolites and the corresponding receptors are significantly affected by MG depletion. Norepinephrine is implicated in both wakefulness^42^ and sleep regulation^43–45^, and it has been suggested that microglial surveillance process relies heavily on norepinephrine signaling^46,47^. Additionally, we observed a decrease in the expression of neurotransmitter precursors such as L-serine, which was mirrored by changes in the concentrations of glycine, asparagine, and glutamic acid. Previous studies have proposed serine as a link between metabolism and neurotransmission^48^, and serine serves as a precursor for synthesizing ceramide ^49^, which has been suggested to be essential for maintaining stable wakefulness ^50^.

Our findings indicate that MG affect vigilane state regulation via multiple pathways and play a more substantial role in maintaining the physical connectivity of brain networks rather than neurochemical connectivity, as MG repopulation largely restores metabolites but not synapses. To gain a detailed understanding of MG regulation of sleep, it will be necessary to explore local specific microglial depletion, and disruption of microglia-neuron interactions using inducible systems. Additionally, investigating the separate contributions of microglia and CAMs in sleep regulation using Cre lines under the promoter of *Hexb* ^51^ and *Mrc1* ^52^ genes, respectively, would be a starting point. The observed vigilance state phenotypic alterations may also be influenced by disruptions in central factors (such as protein phosphorylation orchestrated by microglial TNF-α)^53^, and peripheral players (since CSF1R inhibition also effects circulating and tissue macrophages)^54^ warranting further investigations. Overall, our results shed new light on the complex interaction between the neuroimmune system and sleep regulation, and it paves the way for future investigations in this important physiopathological field.

## Data availability

All data are available from the corresponding author upon reasonable request.

## Acknowledgements

We extend our gratitude to the Metabolomics Core Facility at the Department of General Pediatrics, University of Freiburg, with special thanks to Dr. Luciana Hannibal and Victoria Wingert for their invaluable support and expertise. AS was supported by Novartis Foundation for medical-biological Research (#23B079). MB was supported by the Swiss National Science Foundation (grant 190605). MT was supported by Swiss NSF (grant 201235), and the State of Vaud (Faculty of Biology and Medicine, University of Lausanne). AV was supported by the Swiss National Science Foundation (grant 31003A_182613). AK was supported by institute of neuroinformatics ETH-Zurich. GM was supported by H2020-MSCA-IF grant, n. 101025176.

## Author contributions

AS conceptualized and designed the study. AS, MH, and SW performed experiments. AS, GM, and MB analyzed data. AS, GM, AK, AV, MT, and MB wrote the manuscript.

## Declaration of interests

The authors declare no competing interests.

## Methods

### Animals

Wild type (C57BL/6J) mice were used in this study. All animals were between 12 and 14 weeks old during the time of experiments and animal experiments were approved by the Ministry for Nature, Environment and Consumers’ Protection of the state of Baden-Württemberg and were performed in accordance to the respective national, federal, and institutional regulations. At all times, care was taken to minimize animal discomfort and avoid pain.

### Gene expression analysis

For qPCR analysis, RNA extracted from the cortical samples of control, MG-Dep and MG-Repop mice. RNA was extracted using RNeasy micro kit (74004) and cDNA synthesis was performed using random hexamer primers and M-MLV Reverse Transcriptase Promega kit (M3682). SYBR green-based and Taqman probes were used to detect the gene expression. The assay was performed using a LightCycler480 (Roche).

### Immunofluorescence and synapse counting

Mice were anesthetized with ketamine (100 mg per kg body weight) and xylazine (10 mg per kg body weight) followed by perfusion with 1× PBS. For histology, brains were kept overnight in 4% paraformaldehyde (PFA). Brains were then dehydrated in 30% sucrose for about 48 hours and then snap-frozen on dry ice. Cryosections (14μm for synapse counting and 20μm for microglia and CAM staining) from brain tissue were prepared as free floating in PBS. The following antibodies were used to stain microgliaand synapses: Rabbit-anti-IBA1 (ab178846, abcam), anti-NEUN (ab177487, ab104224, abcam), anti-VLUT1 (135304, synaptic system), anti-HOMER1 (ABN37, Millipore), anti-Gephyrin (147318, synaptic system), anti-Synaptotagmin-2 (ZDB-ATB-081002-25, ZFIN), anti-Parvalbumin (MAB1572, Merck) and anti-HCRT-A (SC-8070). The DAPI was used to stain the nucleus. Iba1 antibody was incubated overnight at 4 °C and the primary antibodies to detect synapses were incubated for 1.45hr at room temperature. The brain cuts were incubated with secondary antibodies for 1-2 hours at room temperature. Fluorescence imaging was performed with TCS SP8 X (Leica Microsystems). Imaging of synapses was performed using 63X objective with zoom factor 8. Iba1^+^ cells were counted using Fiji. Synapse analysis and quantification were conducted using Fiji macro developed previously ^55^ with little modifications.

### Quantitative profiling of metabolites by liquid chromatography and mass spectrometry (LC-MS/MS)

Mice brain tissue biopsies were thawed and homogenised with ice-cold PBS supplemented with 1% protease inhibitor cocktail (Sigma Nr. P8340-5ML), using a cordless pestle motor and disposable pellet mixers (VWR Nr. 47747-366). Buffer volume was adjusted to reach a target of 0.1 to 0.3 mg wet-tissue/mL lysis buffer. Whole tissue homogenates were aliquoted for downstream measurement of total protein concentration, flash-frozen and stored at -80 °C. Sulfur-containing metabolites as well as creatinine, S-adenosylmethionine and S-adenosylhomocysteine were determined according to a previously published procedure ^56,57^. Lactate, TCA and glycolysis intermediates and other organic acids, and folates were determined as described in previous work ^58^. Amino acids and neurotransmitters were determined using a previously described protocol ^59,60^. A commercially available standardized amino acid mixture was utilized to generate a calibration curve for amino acids (Amino acid standards, physiological, Sigma, Nr. A9906-10ML). Calibration curves for all other metabolites were prepared from individual stock solutions prepared in house. Quantitation accuracy was examined by monitoring homocysteine and methylmalonic acid concentrations in an external quality control, namely, the Control Special Assays in Serum, European Research Network for the evaluation and improvement of screening, diagnosis, and treatment of Inherited disorders of Metabolism (ERNDIM) IQCS, SAS-02.1 and SAS-02.2 from MCA Laboratories, Winterswijk, Netherlands. For all other metabolites, quantitation trueness was tested by examining metabolite concentrations in plasma from a previously validated sample isolated from a healthy control individual with respect to standard reference ranges, using the same calibration curves and LC-MS/MS running conditions. Quantification of metabolites was carried out with Analyst® 1.7.2 software, 2022 AB Sciex.

### Treatments

In order to eliminate cells dependent on CSF1R (MG) mice were administered a CSF1R inhibitor called PLX5622 (Plexxikon). The inhibitor was combined with AIN-76A standard chow at a concentration of 1,200ppm (Research Diets) and the mice were given unrestricted access to it for two weeks. The control group received AIN-76A standard chow without any modifications. After two weeks the PLX5622 food was replaced with control food to start repopulation of MG.

### Mice surgery, data acquisition and vigilance state analysis

Mice surgery and data acquisition were performed as previously described ^25^. MATLAB scripts were created to quantify various parameters related to wakefulness, NREMS and REMS. These parameters include time distribution and amount of episodes, as well as the fragmentation of vigilance states, all of which were defined based on previously established criteria ^25^. TDW analysis was conducted following the method described by Vassalli and Franken ^28^.

### Power spectral analysis

Using Somnologica-3TM (Medcare) software, each 4 s epoch of the EEG signal was subjected to discrete Fourier transform to determine EEG power density (spectra 0–90 Hz at a frequency resolution of 0.25 Hz). We used artifact-free, same-state–flanked 4 s epochs to calculate the mean EEG spectral profiles for each behavioral states and time intervals. To account for differences among animals in absolute EEG power, the mean values of spectral profiles were converted into percentages relative to a baseline EEG power reference value of 100%. This reference value was determined for each mouse by summing the power within the frequency range of 0.75–47.5 Hz across all states and throughout the first day of baseline recordings. To control for differences in the amount of wakefulness, NREMS and REMS, this reference was weighted so that the relative contribution of each state was identical for all mice ^61^. Time-frequency heatmaps of EEG power during wakefulness were generated using methods described previously ^62^.

### Time-course analysis

To examine the dynamic changes of EEG power within specific frequency ranges throughout the day and night, the time-course of the activity of that frequency band was computed for 4 s epochs scored as state of interest (NREMS, REMS and wake). For this purpose, the epochs for each state were divided into percentiles, ensuring that each percentile contained a roughly equal number of epochs. The number of percentiles for NREMS and REMS were: 12 for light periods, 6 for dark periods, and 8 for the 6-h light period after SD; for waking state: 6 for light periods, 12 for dark periods, 8 for the 6-h light SD, and 4 for the 6-h light after SD. The average EEG power within the specified frequency range was computed for each percentile, and subsequently normalized based on the type of analysis. This normalization was either done relative to the total power of the baseline state or with respect to the power achieved within the delta frequency band during slow-wave sleep at ZT8-12. Like power density analysis, single epochs were excluded and only power values of the epochs that themselves, as well as the two adjacent ones, were scored as artifact-free same-state were included in the analysis.

### Theta-gamma cross-frequency coupling

We used the modulation index (MI) to measure the theta-gamma phase-amplitude coupling ^27,63^. Using finite impulse response filters with an order equal to three cycles of the low cutoff frequency, we first bandpass-filtered EEG signals into theta (6-11 Hz) and fast-gamma (54-90 Hz) in both forward and reverse directions to eliminate phase distortion. We then estimated instantaneous phase of theta and the envelope of fast-gamma using the Hilbert transform. Theta phase was discretized into 18 equal bins (*N*=18, each 20°) and the average value of fast-gamma envelope within each bin was calculated. The resulting phase-amplitude histogram (*P*) was compared with a uniform distribution (*U*) using the Kullback-Leibler distance, *D_KL_*(*P, U*) = Σ^*N*^_*j*=1_*P*(*j*)* *log*[*P*(*j*)/*U*(*j*)], and normalized by *log(N)* to obtain the modulation index, *MI = D_KL_ / log(N)*. To explore possible coupling patterns between different pairs of low and high frequency bands, we used the comodulogram analysis ^63^. We considered 16 frequency bands for phase (1-18 Hz, 1-Hz increments, 2-Hz bandwidth), and 14 frequency bands for amplitude (15-90 Hz, 5-Hz increments, 10-Hz bandwidth). MI values were then calculated for all these pairs to obtain the comodulogram graph.

### Detection of sleep spindles

NREMS spindles were detected automatically using an optimized wavelet-based method as previously described ^26^. Briefly, the power of EEG signals within 9-16Hz was estimated using the complex B-spline wavelet function, and smoothed using a 200 ms Hanning window, and then a threshold equal to 3 SD (SD: standard deviation) above the mean was applied to detect the potential spindle events. Events shorter than 400 ms or longer than 2 s were discarded. Using band pass-filtered EEG signals in the spindle range (9–16 Hz), we automatically counted the number of cycles of each detected event and excluded those with <5 cycles or more than 30 cycles. To discard artefacts, events with a power in the spindle band lower than 6–8.5 Hz or 16.5–20 Hz power bands were not included.

### Statistical analysis

The specific number (n) of replicates employed in each experiment is provided in the corresponding figure legends. The individuals responsible for conducting SD and scoring the sleep recordings were unaware of the genotype of the animals, ensuring blinding during these procedures. The results are presented as mean ± SEM or mean ± SD. Statistical analyses were conducted using GraphPad Prism. The significance of comparisons was determined using one-or two-way ANOVA or mix effect model followed by relevant post hoc tests and are reported in the figure legends.

## Notes

### Competing Interest Statement

The authors have declared no competing interest.

### Summary of Updates

The entire story has been improved and updated. Author list is updated.

## References

1. Ramar, K., Malhotra, R.K., Carden, K.A., Martin, J.L., Abbasi-Feinberg, F., Aurora, R.N., Kapur, V.K., Olson, E.J., Rosen, C.L., Rowley, J.A., et al. (2021). Sleep is essential to health: an American Academy of Sleep Medicine position statement. J Clin Sleep Med. 10.5664/jcsm.9476.

2. Borbély, A.A. (1982). A two process model of sleep regulation. Hum Neurobiol 1, 195–204.

3. Gilestro, G.F., Tononi, G., and Cirelli, C. (2009). Widespread changes in synaptic markers as a function of sleep and wakefulness in Drosophila. Science 324, 109–112. 10.1126/science.1166673.

4. Maret, S., Dorsaz, S., Gurcel, L., Pradervand, S., Petit, B., Pfister, C., Hagenbuchle, O., O’Hara, B.F., Franken, P., and Tafti, M. (2007). Homer1a is a core brain molecular correlate of sleep loss. Proc Natl Acad Sci U S A 104, 20090–20095. 10.1073/pnas.0710131104.

5. Saper, C.B., Chou, T.C., and Scammell, T.E. (2001). The sleep switch: hypothalamic control of sleep and wakefulness. Trends Neurosci 24, 726–731. 10.1016/s0166-2236(00)02002-6.

6. Amann, L., Masuda, T., and Prinz, M. (2023). Mechanisms of myeloid cell entry to the healthy and diseased central nervous system. Nat Immunol 24, 393–407. 10.1038/s41590-022-01415-8.

7. Kierdorf, K., Masuda, T., Jordão, M.J.C., and Prinz, M. (2019). Macrophages at CNS interfaces: ontogeny and function in health and disease. Nat Rev Neurosci 20, 547–562. 10.1038/s41583-019-0201-x.

8. Prinz, M., Masuda, T., Wheeler, M.A., and Quintana, F.J. (2021). Microglia and Central Nervous System-Associated Macrophages-From Origin to Disease Modulation. Annual review of immunology 39, 251–277. 10.1146/annurev-immunol-093019-110159.

9. Prinz, M., Jung, S., and Priller, J. (2019). Microglia Biology: One Century of Evolving Concepts. Cell 179, 292–311. 10.1016/j.cell.2019.08.053.

10. Paolicelli, R.C., Bolasco, G., Pagani, F., Maggi, L., Scianni, M., Panzanelli, P., Giustetto, M., Ferreira, T.A., Guiducci, E., Dumas, L., et al. (2011). Synaptic pruning by microglia is necessary for normal brain development. Science (New York, N.Y.) 333, 1456–1458. 10.1126/science.1202529.

11. Shigemoto-Mogami, Y., Hoshikawa, K., Goldman, J.E., Sekino, Y., and Sato, K. (2014). Microglia enhance neurogenesis and oligodendrogenesis in the early postnatal subventricular zone. J Neurosci 34, 2231–2243. 10.1523/jneurosci.1619-13.2014.

12. Nimmerjahn, A., Kirchhoff, F., and Helmchen, F. (2005). Resting microglial cells are highly dynamic surveillants of brain parenchyma in vivo. Science 308, 1314–1318. 10.1126/science.1110647.

13. Kierdorf, K., Erny, D., Goldmann, T., Sander, V., Schulz, C., Perdiguero, E.G., Wieghofer, P., Heinrich, A., Riemke, P., Hölscher, C., et al. (2013). Microglia emerge from erythromyeloid precursors via Pu.1- and Irf8-dependent pathways. Nature neuroscience 16, 273–280. 10.1038/nn.3318.

14. Schulz, C., Gomez Perdiguero, E., Chorro, L., Szabo-Rogers, H., Cagnard, N., Kierdorf, K., Prinz, M., Wu, B., Jacobsen, S.E., Pollard, J.W., et al. (2012). A lineage of myeloid cells independent of Myb and hematopoietic stem cells. Science (New York, N.Y.) 336, 86–90. 10.1126/science.1219179.

15. Bruttger, J., Karram, K., Wörtge, S., Regen, T., Marini, F., Hoppmann, N., Klein, M., Blank, T., Yona, S., Wolf, Y., et al. (2015). Genetic Cell Ablation Reveals Clusters of Local Self-Renewing Microglia in the Mammalian Central Nervous System. Immunity 43, 92–106. 10.1016/j.immuni.2015.06.012.

16. Tay, T.L., Mai, D., Dautzenberg, J., Fernández-Klett, F., Lin, G., Sagar, Datta M., Drougard, A., Stempfl, T., Ardura-Fabregat, A., et al. (2017). A new fate mapping system reveals context-dependent random or clonal expansion of microglia. Nature neuroscience 20, 793–803. 10.1038/nn.4547.

17. Huang, Y., Xu, Z., Xiong, S., Sun, F., Qin, G., Hu, G., Wang, J., Zhao, L., Liang, Y.X., Wu, T., et al. (2018). Repopulated microglia are solely derived from the proliferation of residual microglia after acute depletion. Nat Neurosci 21, 530–540. 10.1038/s41593-018-0090-8.

18. Ingiosi, A.M., Opp, M.R., and Krueger, J.M. (2013). Sleep and immune function: glial contributions and consequences of aging. Curr Opin Neurobiol 23, 806–811. 10.1016/j.conb.2013.02.003.

19. Imeri, L., and Opp, M.R. (2009). How (and why) the immune system makes us sleep. Nat Rev Neurosci 10, 199–210. 10.1038/nrn2576.

20. Tuan, L.H., and Lee, L.J. (2019). Microglia-mediated synaptic pruning is impaired in sleep-deprived adolescent mice. Neurobiol Dis 130, 104517. 10.1016/j.nbd.2019.104517.

21. Gentry, N.W., McMahon, T., Yamazaki, M., Webb, J., Arnold, T.D., Rosi, S., Ptáček, L.J., and Fu, Y.H. (2022). Microglia are involved in the protection of memories formed during sleep deprivation. Neurobiol Sleep Circadian Rhythms 12, 100073. 10.1016/j.nbscr.2021.100073.

22. Bellesi, M., de Vivo, L., Chini, M., Gilli, F., Tononi, G., and Cirelli, C. (2017). Sleep Loss Promotes Astrocytic Phagocytosis and Microglial Activation in Mouse Cerebral Cortex. J Neurosci 37, 5263–5273. 10.1523/jneurosci.3981-16.2017.

23. Dagher, N.N., Najafi, A.R., Kayala, K.M., Elmore, M.R., White, T.E., Medeiros, R., West, B.L., and Green, K.N. (2015). Colony-stimulating factor 1 receptor inhibition prevents microglial plaque association and improves cognition in 3xTg-AD mice. J Neuroinflammation 12, 139. 10.1186/s12974-015-0366-9.

24. Van Hove, H., Martens, L., Scheyltjens, I., De Vlaminck, K., Pombo Antunes, A.R., De Prijck, S., Vandamme, N., De Schepper, S., Van Isterdael, G., Scott, C.L., et al. (2019). A single-cell atlas of mouse brain macrophages reveals unique transcriptional identities shaped by ontogeny and tissue environment. Nat Neurosci 22, 1021–1035. 10.1038/s41593-019-0393-4.

25. Seifinejad, A., Li, S., Possovre, M.L., Vassalli, A., and Tafti, M. (2020). Hypocretinergic interactions with the serotonergic system regulate REM sleep and cataplexy. Nat Commun 11, 6034. 10.1038/s41467-020-19862-y.

26. Bandarabadi, M., Herrera, C.G., Gent, T.C., Bassetti, C., Schindler, K., and Adamantidis, A.R. (2020). A role for spindles in the onset of rapid eye movement sleep. Nat Commun 11, 5247. 10.1038/s41467-020-19076-2.

27. Bandarabadi, M., Boyce, R., Gutierrez Herrera, C., Bassetti, C.L., Williams, S., Schindler, K., and Adamantidis, A. (2019). Dynamic modulation of theta-gamma coupling during rapid eye movement sleep. Sleep 42, 1–11. 10.1093/sleep/zsz182.

28. Vassalli, A., and Franken, P. (2017). Hypocretin (orexin) is critical in sustaining theta/gamma-rich waking behaviors that drive sleep need. Proc Natl Acad Sci U S A 114, E5464–e5473. 10.1073/pnas.1700983114.

29. Endo, T., Roth, C., Landolt, H.P., Werth, E., Aeschbach, D., Achermann, P., and Borbély, A.A. (1998). Selective REM sleep deprivation in humans: effects on sleep and sleep EEG. Am J Physiol 274, R1186–1194. 10.1152/ajpregu.1998.274.4.R1186.

30. Tononi, G., and Cirelli, C. (2003). Sleep and synaptic homeostasis: a hypothesis. Brain Res Bull 62, 143–150. 10.1016/j.brainresbull.2003.09.004.

31. Hristovska, I., Robert, M., Combet, K., Honnorat, J., Comte, J.C., and Pascual, O. (2022). Sleep decreases neuronal activity control of microglial dynamics in mice. Nat Commun 13, 6273. 10.1038/s41467-022-34035-9.

32. Kon, K., Ode, K.L., Mano, T., Fujishima, H., Tone, D., Shimizu, C., Shiono, S., Yada, S., Garçon, J.Y., Kaneko, M., et al. (2023). Cortical parvalbumin neurons are responsible for homeostatic sleep rebound through CaMKII activation. bioRxiv, 2023.2004.2029.537929. 10.1101/2023.04.29.537929.

33. Bouhours, B., Gjoni, E., Kochubey, O., and Schneggenburger, R. (2017). Synaptotagmin2 (Syt2) Drives Fast Release Redundantly with Syt1 at the Output Synapses of Parvalbumin-Expressing Inhibitory Neurons. J Neurosci 37, 4604–4617. 10.1523/jneurosci.3736-16.2017.

34. Morairty, S.R., Dittrich, L., Pasumarthi, R.K., Valladao, D., Heiss, J.E., Gerashchenko, D., and Kilduff, T.S. (2013). A role for cortical nNOS/NK1 neurons in coupling homeostatic sleep drive to EEG slow wave activity. Proc Natl Acad Sci U S A 110, 20272–20277. 10.1073/pnas.1314762110.

35. Cirelli, C., Huber, R., Gopalakrishnan, A., Southard, T.L., and Tononi, G. (2005). Locus ceruleus control of slow-wave homeostasis. J Neurosci 25, 4503–4511. 10.1523/jneurosci.4845-04.2005.

36. Willis, E.F., MacDonald, K.P.A., Nguyen, Q.H., Garrido, A.L., Gillespie, E.R., Harley, S.B.R., Bartlett, P.F., Schroder, W.A., Yates, A.G., Anthony, D.C., et al. (2020). Repopulating Microglia Promote Brain Repair in an IL-6-Dependent Manner. Cell 180, 833–846.e816. 10.1016/j.cell.2020.02.013.

37. Zhang, Y., Zhao, L., Wang, X., Ma, W., Lazere, A., Qian, H.H., Zhang, J., Abu-Asab, M., Fariss, R.N., Roger, J.E., and Wong, W.T. (2018). Repopulating retinal microglia restore endogenous organization and function under CX3CL1-CX3CR1 regulation. Sci Adv 4, eaap8492. 10.1126/sciadv.aap8492.

38. Favuzzi, E., Huang, S., Saldi, G.A., Binan, L., Ibrahim, L.A., Fernández-Otero, M., Cao, Y., Zeine, A., Sefah, A., Zheng, K., et al. (2021). GABA-receptive microglia selectively sculpt developing inhibitory circuits. Cell 184, 4048–4063.e4032. 10.1016/j.cell.2021.06.018.

39. Wilton, D.K., Dissing-Olesen, L., and Stevens, B. (2019). Neuron-Glia Signaling in Synapse Elimination. Annu Rev Neurosci 42, 107–127. 10.1146/annurev-neuro-070918-050306.

40. Badimon, A., Strasburger, H.J., Ayata, P., Chen, X., Nair, A., Ikegami, A., Hwang, P., Chan, A.T., Graves, S.M., Uweru, J.O., et al. (2020). Negative feedback control of neuronal activity by microglia. Nature 586, 417–423. 10.1038/s41586-020-2777-8.

41. Guillaumin, M.C.C., Harding, C.D., Krone, L.B., Yamagata, T., Kahn, M.C., Blanco-Duque, C., Banks, G.T., Nolan, P.M., Peirson, S.N., and Vyazovskiy, V.V. (2023). Deficient synaptic neurotransmission results in a persistent sleep-like cortical activity across vigilance states in mice. bioRxiv, 2023.2005.2011.540034. 10.1101/2023.05.11.540034.

42. Seifinejad, A., Vassalli, A., and Tafti, M. (2021). Neurobiology of cataplexy. Sleep Med Rev 60, 101546. 10.1016/j.smrv.2021.101546.

43. Osorio-Forero, A., Cardis, R., Vantomme, G., Guillaume-Gentil, A., Katsioudi, G., Devenoges, C., Fernandez, L.M.J., and Lüthi, A. (2021). Noradrenergic circuit control of non-REM sleep substates. Current biology : CB 31, 5009–5023.e5007. 10.1016/j.cub.2021.09.041.

44. Ma, C., Li, B., Silverman, D., Ding, X., Li, A., Xiao, C., Huang, G., Worden, K., Muroy, S., Chen, W., et al. (2023). Microglia Regulate Sleep via Calcium-Dependent Modulation of Norepinephrine Transmission. bioRxiv, 2023.2007.2024.550176. 10.1101/2023.07.24.550176.

45. Ma, C., Li, B., Silverman, D., Ding, X., Li, A., Xiao, C., Huang, G., Worden, K., Muroy, S., Chen, W., et al. (2024). Microglia regulate sleep through calcium-dependent modulation of norepinephrine transmission. Nat Neurosci 27, 249–258. 10.1038/s41593-023-01548-5.

46. Stowell, R.D., Sipe, G.O., Dawes, R.P., Batchelor, H.N., Lordy, K.A., Whitelaw, B.S., Stoessel, M.B., Bidlack, J.M., Brown, E., Sur, M., and Majewska, A.K. (2019). Noradrenergic signaling in the wakeful state inhibits microglial surveillance and synaptic plasticity in the mouse visual cortex. Nat Neurosci 22, 1782–1792. 10.1038/s41593-019-0514-0.

47. Liu, Y.U., Ying, Y., Li, Y., Eyo, U.B., Chen, T., Zheng, J., Umpierre, A.D., Zhu, J., Bosco, D.B., Dong, H., and Wu, L.J. (2019). Neuronal network activity controls microglial process surveillance in awake mice via norepinephrine signaling. Nat Neurosci 22, 1771–1781. 10.1038/s41593-019-0511-3.

48. Maugard, M., Vigneron, P.A., Bolanos, J.P., and Bonvento, G. (2021). l-Serine links metabolism with neurotransmission. Prog Neurobiol 197, 101896. 10.1016/j.pneurobio.2020.101896.

49. Vávrová, K., Hrabálek, A., Dolezal, P., Holas, T., and Zbytovská, J. (2003). L-Serine and glycine based ceramide analogues as transdermal permeation enhancers: polar head size and hydrogen bonding. Bioorg Med Chem Lett 13, 2351–2353. 10.1016/s0960-894x(03)00409-8.

50. Liu, H., Wang, X., Chen, L., Chen, L., Tsirka, S.E., Ge, S., and Xiong, Q. (2021). Microglia modulate stable wakefulness via the thalamic reticular nucleus in mice. Nat Commun 12, 4646. 10.1038/s41467-021-24915-x.

51. Masuda, T., Amann, L., Sankowski, R., Staszewski, O., Lenz, M.P.D.E., Snaidero, N., Costa Jordão, M.J., Böttcher, C., Kierdorf, K., et al. (2020). Novel Hexb-based tools for studying microglia in the CNS. Nat Immunol 21, 802–815. 10.1038/s41590-020-0707-4.

52. Masuda, T., Amann, L., Monaco, G., Sankowski, R., Staszewski, O., Krueger, M., Del Gaudio, F., He, L., Paterson, N., Nent, E., et al. (2022). Specification of CNS macrophage subsets occurs postnatally in defined niches. Nature 604, 740–748. 10.1038/s41586-022-04596-2.

53. Pinto, M.J., Cottin, L., Dingli, F., Laigle, V., Ribeiro, L.F., Triller, A., Henderson, F., Loew, D., Fabre, V., and Bessis, A. (2023). Microglial TNFα orchestrates protein phosphorylation in the cortex during the sleep period and controls homeostatic sleep. Embo j 42, e111485. 10.15252/embj.2022111485.

54. Lei, F., Cui, N., Zhou, C., Chodosh, J., Vavvas, D.G., and Paschalis, E.I. (2020). CSF1R inhibition by a small-molecule inhibitor is not microglia specific; affecting hematopoiesis and the function of macrophages. Proc Natl Acad Sci U S A 117, 23336–23338. 10.1073/pnas.1922788117.

55. Favuzzi, E., Huang, S., Saldi, G.A., Binan, L., Ibrahim, L.A., Fernández-Otero, M., Cao, Y., Zeine, A., Sefah, A., Zheng, K., et al. (2021). GABA-receptive microglia selectively sculpt developing inhibitory circuits. Cell 184, 5686. 10.1016/j.cell.2021.10.009.

56. Behringer, S., Wingert, V., Oria, V., Schumann, A., Grunert, S., Cieslar-Pobuda, A., Kolker, S., Lederer, A.K., Jacobsen, D.W., Staerk, J., et al. (2019). Targeted Metabolic Profiling of Methionine Cycle Metabolites and Redox Thiol Pools in Mammalian Plasma, Cells and Urine. Metabolites 9. 10.3390/metabo9100235.

57. Bravo, A.C., Aguilera, M.N.L., Marziali, N.R., Moritz, L., Wingert, V., Klotz, K., Schumann, A., Grünert, S.C., Spiekerkoetter, U., Berger, U., et al. (2022). Analysis of S-Adenosylmethionine and S-Adenosylhomocysteine: Method Optimisation and Profiling in Healthy Adults upon Short-Term Dietary Intervention. Metabolites 12, 373.

58. Hannibal, L., Theimer, J., Wingert, V., Klotz, K., Bierschenk, I., Nitschke, R., Spiekerkoetter, U., and Grunert, S.C. (2020). Metabolic Profiling in Human Fibroblasts Enables Subtype Clustering in Glycogen Storage Disease. Front Endocrinol (Lausanne) 11, 579981. 10.3389/fendo.2020.579981.

59. Maier, J.P., Ravi, V.M., Kueckelhaus, J., Behringer, S.P., Garrelfs, N., Will, P., Sun, N., von Ehr, J., Goeldner, J.M., Pfeifer, D., et al. (2021). Inhibition of metabotropic glutamate receptor III facilitates sensitization to alkylating chemotherapeutics in glioblastoma. Cell Death Dis 12, 723. 10.1038/s41419-021-03937-9.

60. Moritz, L., Klotz, K., Grunert, S.C., Hannibal, L., and Spiekerkoetter, U. (2023). Metabolic phenotyping in phenylketonuria reveals disease clustering independently of metabolic control. Mol Genet Metab 138, 107509. 10.1016/j.ymgme.2023.107509.

61. Franken, P., Malafosse, A., and Tafti, M. (1998). Genetic variation in EEG activity during sleep in inbred mice. Am J Physiol 275, R1127–1137. 10.1152/ajpregu.1998.275.4.R1127.

62. Li, S., Franken, P., and Vassalli, A. (2018). Bidirectional and context-dependent changes in theta and gamma oscillatory brain activity in noradrenergic cell-specific Hypocretin/Orexin receptor 1-KO mice. Sci Rep 8, 15474. 10.1038/s41598-018-33069-8.

63. Tort, A.B., Kramer, M.A., Thorn, C., Gibson, D.J., Kubota, Y., Graybiel, A.M., and Kopell, N.J. (2008). Dynamic cross-frequency couplings of local field potential oscillations in rat striatum and hippocampus during performance of a T-maze task. Proc Natl Acad Sci U S A 105, 20517–20522. 10.1073/pnas.0810524105.

